# Elucidating a potential role of the infant gut microbiome on the bioavailability of L-tyrosine in phenylketonuria

**DOI:** 10.1101/2025.10.28.685037

**Authors:** Mohammadreza Moghimi, Filippo Martinelli, Jonas Widder, Tim Hensen, Luciana Hannibal, Ute Spiekerkoetter, Ines Thiele

## Abstract

**Background:** Phenylketonuria (PKU) is an inherited metabolic disorder caused by phenylalanine hydroxylase (PAH) deficiency, leading to elevated L-phenylalanine and severe neurological damage if untreated. While phenylalanine-based biomarkers are diagnostic and phenylalanine levels correlate with disease severity, the clinical manifestations of PKU are heterogeneous.

**Results:** To identify additional reliable, potentially novel biomarkers, we used germ-free sex-specific, organ-resolved infant whole-body metabolic models (infant-WBMs) to simulate PAH deficiency and predicted elevated L-phenylalanine and its derivatives, alongside reduced L-tyrosine fluxes, as the product of phenylalanine hydroxylation. To test the reliability of these predictions, we combined the infant-WBMs with gut microbiome models from 48 healthy infants. Upon integrating microbiome data, we found that microbial metabolism significantly increased L-tyrosine availability, obscuring its utility as a universal biomarker. In ∼23% of microbiome-PKU models, L-tyrosine fluxes remained low, indicating insufficient microbial compensation. These cases were enriched in Firmicutes and lacked specific Bifidobacterium and Escherichia strains linked to L-tyrosine biosynthesis via the pretyrosine pathway. Shadow price analysis identified microbial species critical for host L-tyrosine metabolism. However, some, such as *Bifidobacterium dentium*, also contributed to L-phenylalanine synthesis, potentially worsening the PKU phenotype. In contrast, L-phenylalanine, phenylpyruvate, and hydroxyphenylacetic acid remained reliably elevated across all models, validating their diagnostic relevance.

**Conclusions:** Our study demonstrates that microbiome composition can modulate biomarker reliability in PKU, particularly for L-tyrosine. Integrating microbial metabolic models with whole-body physiology enables assessment of biomarker reliability and reveals subpopulations for whom secondary biomarkers or targeted probiotics may be beneficial. This approach offers a powerful framework for refining diagnostics and therapy monitoring in rare metabolic diseases and the development of possible targeted microbiome therapies.

## Introduction

Phenylketonuria (PKU) is a rare genetic disorder caused by a deficiency in phenylalanine hydroxylase (PAH), the enzyme responsible for converting L-phenylalanine into L-tyrosine with tetrahydrobiopterin (BH₄) as a cofactor ^1^. Newborn-screening programmes test for PKU by measuring the L-phenylalanine levels in dried-blood spots taken from neonates (between 24 to 72 hours after birth). Early detection of PAH deficiency is paramount as L-phenylalanine accumulates in the brain and blood, which leads to non-reversible neurological damage ^2^. In addition, L-tyrosine levels are reduced in some but not all PKU patients ^3^. Since L-tyrosine is a precursor for synthesis of neurotransmitters, such as dopamine, norepinephrine, epinephrine, and serotonin ^3^, as well as melanin and thyroid hormones, a decreased availability of L-tyrosine contributes to cognitive impairments, mood disturbances, attention deficits, lighter pigmentation, and metabolic disruptions ^4–6^. Without treatment, PKU leads to intellectual disability and severe developmental delays ^7^. However, early diagnosis through newborn screening and strict dietary management prevents these complications ^5^. PKU treatment focuses on controlling blood L-phenylalanine levels to prevent toxicity, while ensuring sufficient L-tyrosine and other essential amino acids for cognitive and metabolic functions ^8,9^. The primary approach is a strict low-L-phenylalanine diet that excludes high-protein foods, such as meat, fish, dairy, and nuts, replaced with specially formulated medical foods, which contain synthetic protein to provide all essential amino acids in the necessary amounts ^9,10^. While effective, this diet often results in low L-tyrosine levels, necessitating supplementation ^1,10^. Additional therapeutic strategies include BH₄ supplementation, enzyme replacement, and gene therapy approaches ^1,5^.

L-phenylalanine in blood and cerebrospinal fluid, as well as phenylpyruvic acid in urine, are established biomarkers for the diagnosis of PKU ^11^. Despite the significance of L-phenylalanine in the diagnosis of PKU, and the correlation of high L-phenylalanine blood levels with disease severity, the heterogeneous phenotypes with respect to the degree of cognitive impairment are not completely understood ^12^ ^13^. Hence, there is a need for novel biomarkers that can be further linked to the heterogenous phenotypes. Other metabolites, which have been reported to be altered in PKU patients include hydroxyphenylacetic acid ^12^ and L-tyrosine ^14^. However, L-tyrosine levels have been reported to be low to normal in PKU patients ^3^ and thus, its use as a biomarker remains unclear. Additionally, phenylalanine and L-tyrosine levels do fluctuate in patients during the day depending on food and aminoacid mixure intake.

The suitability of diagnostic and prognostic biomarkers depends on numerous factors, including the patient’s genetics, diet, and gut microbiome. For instance, the gut microbiota can influence the metabolism of L-phenylalanine and L-tyrosine ^15^, as certain bacterial strains, such as *Bacteroides thetaiotaomicron*, *Bacteroides eggerthii*, and *Bacteroides ovatus*, can degrade L-phenylalanine or convert it into less harmful metabolites ^16^. Thus, their presence in the microbiome could lead to higher tyrosine availability ^17^. However, only a few studies involving PKU patients have investigated microbiome changes associated with the disease and the dietary treatment ^17,18^. Additionally, it has been shown that the diet can alter the microbial composition ^19^. In particular, the restrictive PKU diet with the exclusion of natural protein- and fibre-rich foods, reduces microbial diversity compared to healthy individuals ^17,18^. These microbial changes could affect digestion, gut health, and metabolic processes ^20^. Hence, microbial metabolic activity should be considered in the discovery of novel candidate biomarkers.

Various computational modelling approaches have been used to predict known and novel biomarkers for rare metabolic diseases ^21,22^. One of the most prominent modelling approaches is the constraint-based reconstruction and analysis (COBRA) approach. In COBRA, a genome-scale metabolic reconstruction is built based on an organism’s genome annotation, biochemical, and physiological data ^23^. As such, a metabolic reconstruction describes the biochemical relationships between metabolites in a stoichiometrically accurate manner. Metabolic reconstructions can be tailored to a person or a condition (i.e., a specific diet) by applying limits (i.e., constraints) to reaction fluxes, thereby converting a generic reconstruction to a person- and condition-specific metabolic model. Importantly, and in distinction to other metabolic network approaches, the emerging phenotypes can be predicted with metabolic modelling ^23^, with flux balance analysis being the most widely used method ^24^. While metabolic models are metabolite- and reaction-centric, the enzymes catalysing the reactions and their genes are captured as Boolean rules (‘AND’ or ‘OR’). Consequently, any change in gene activity, e.g., due to a pathogenic genetic variant, can be modelled by adjusting the constraints on the corresponding reaction fluxes. This approach has been applied to predict known and novel candidate metabolite biomarkers from known gene defects ^25–28^.

A recent advance in metabolic modelling has been the development of sex-specific, organ-resolved whole-body models (WBMs) of human metabolism ^28^. These WBMs account for >80,000 biochemical reactions and >60,000 metabolites across 26 organs, six blood cell types, and 13 biofluids. Importantly, the WBMs can be personalised using dietary, metabolomic, physiological, and microbiome data ^28^. These comprehensive models enable studying metabolic interactions between different organs and provide insight into systemic metabolism ^29,30^. WBMs allow researchers to simulate normal human physiology as well as pathophysiological conditions, making them particularly useful in exploring the complexities of inherited metabolic diseases (IMDs) ^28^. In fact, WBMs can predict known biomarker metabolites for over 50 different IMDs ^28^. Equally, these WBMs could be used to suggest novel diagnostic biomarkers.

WBMs have been followed by the development of sex-specific infant whole-body models (infant-WBMs), which are specifically designed to simulate the metabolism of newborns and infants ^31^. These infant-WBMs address the unique metabolic and physiological requirements of early human life, which differ significantly from those of adults. Infant-WBMs extend the capabilities of adult WBMs by incorporating the unique metabolic needs of newborns, such as energy demands for rapid growth and brain development. By incorporating parameters, such as age-dependent organ weights, blood flow rates, and nutritional intake, the infant WBMs provide a comprehensive tool for understanding infant metabolism at an organ-resolved level. They also allow for the prediction of growth trajectories and changes in metabolic biomarkers, including those related to IMDs. Personalised with data from newborn screenings, such as blood metabolite concentrations and birth weight, these infant-WBMs can predict how IMDs, such as PKU, may develop and affect infants, offering valuable insights into early diagnosis and treatment planning ^31^.

Another unique feature of WBMs is that they can be integrated with microbiome metabolic models, generated based on metagenomic data; thereby, allowing for the simulation of host-microbiome co-metabolism ^28^. This integration requires the availability of metabolic reconstructions for microbial strains, or species, identified in a given microbiome sample. To this end, two large-scale resources of microbial metabolic reconstructions are available, being AGORA2 ^32^, which account for 7,302 human gut microbial strains, and APOLLO ^33^, which accounts for 247,092 metagenome assembled genomes microbial metabolic reconstructions and microbial community models for 14,451 metagenomes from multiple body sites. Hence, these resources are well suited for investigating the effect of microbiome metabolic activity on the reliability of predicted known and novel biomarkers in rare metabolic diseases.

## Result

In this study, we aimed to identify reliable biomarkers for PKU. Therefore, we performed metabolic modelling with sex-specific, organ-resolved infant whole-body metabolic models (infant-WBMs). First, simulating PAH deficiency in germ-free infant-WBMs predicted elevated L-phenylalanine and its derivatives, alongside reduced L-tyrosine fluxes. Subsequently, we integrated microbial community models with the infant-WBMs and found that microbial metabolism significantly increased L-tyrosine availability, except for a subset of microbiomes. These cases were enriched in Firmicutes and lacked specific Bifidobacterium and Escherichia strains, which we could link to changes in L-tyrosine biosynthesis via the pretyrosine pathway. Taken together, our approach demonstrates its potential for the identification of reliable biomarkers as well as biomarkers suitable for a subcohort of individuals.

### Prediction of candidate biomarkers for PKU

First, we investigated whether the infant-WBMs could be used to predict novel diagnostic biomarkers. As the newborn screening programmes in most countries are carried out between 24 and 72 hours after birth, we decided to use the female (germ-free) infant-WBM corresponding to day 2. We simulated PKU in the infant-WBMs by setting the bounds of the PAH gene associated reactions associated, i.e., PHETHPTOX2 and r0399, to zero in all organs they appeared in (Methods). Using a pre-selected set of 233 metabolites, which was assembled from biomarkers used in newborn screening programmes and reported IMD biomarkers in IEMBase ^34^, we tested the healthy and PKU infant-WBMs for the maximal production capability of each metabolite in the blood compartment. As expected, L-phenylalanine (Virtual Metabolic Human, VMH ID: phe_L), phenylpyruvate (VMH ID: phpyr), and ortho-hydroxylacetic acid (VMH ID: 2hyoxplac) were increased in the PKU infant-WBMs (**Figure 1**, **Table S1**). The PKU infant-WBM had a reduced flux for L-tyrosine (VMH ID: tyr_L) and its derivatives compared to the healthy infant-WBM (**Figure 1**). None of the other tested metabolites increased or decreased to the same extend as the aforementioned metabolites. However, some small reductions were predicted for a further 61 metabolites, many of which were amino acids (**Table S1**). Notably, while L-phenylalanine and phenylpyruvate are known biomarkers for PKU ^34^, the seven other metabolites have not yet been established, and thus, are candidate novel biomarkers. However, our predicted decreased L-tyrosine availability in the blood compartment is partially inconsistent with reports of serum L-tyrosine levels being low to normal in PKU patients ^3^.

**Figure 1.**
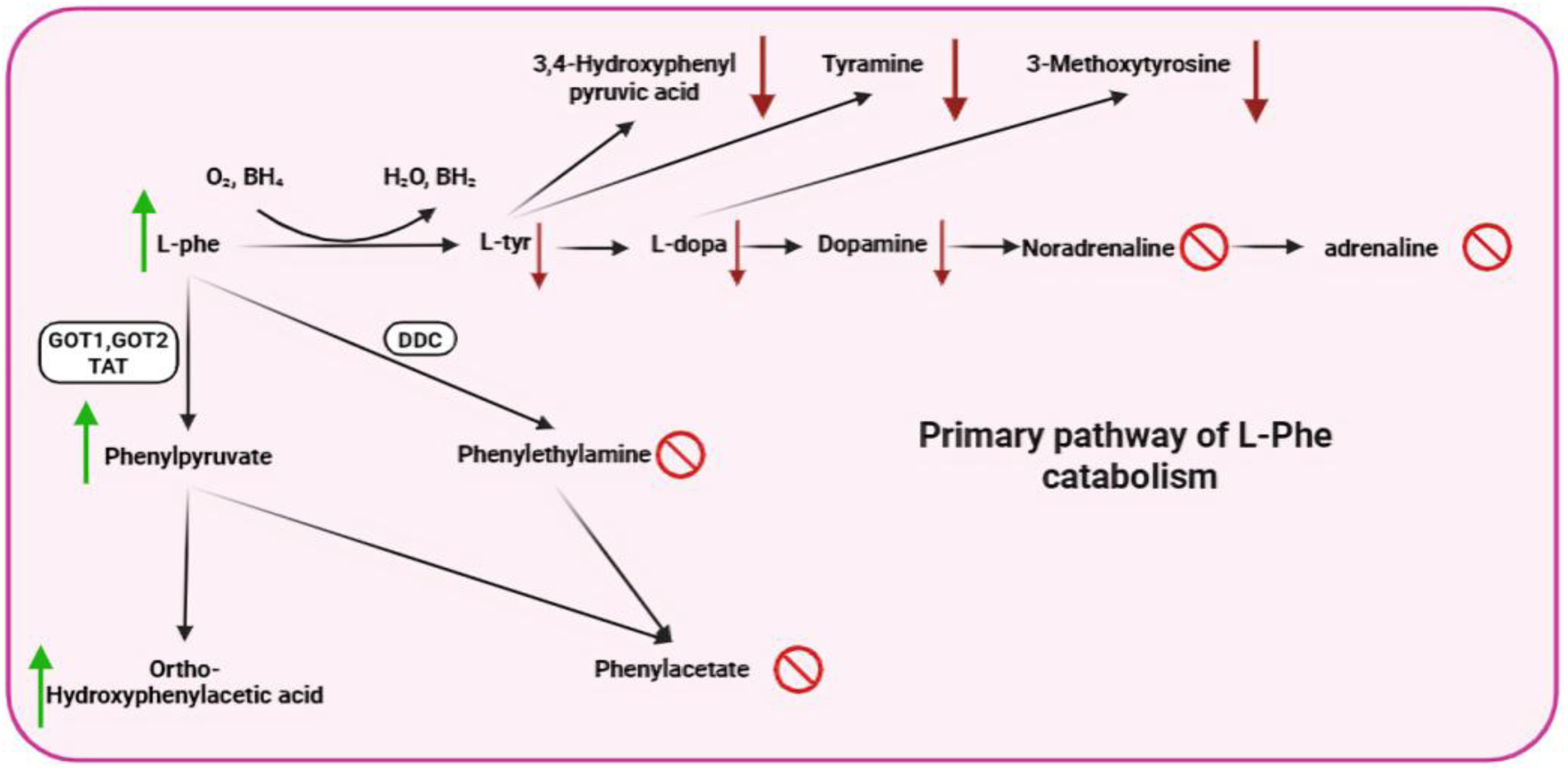
Schematic representation of possible PKU biomarkers in the L-phenylalanine and L-tyrosine metabolism. Metabolites accompanied by arrows indicate biomarkers investigated in this study, with the arrow direction representing their elevation or decrease in PKU. Metabolites accompanied by the prohibition symbol represent potential key biomarkers not investigated in this study. L-phe = L-phenylalanine. L-tyr = L-tyrosine. L-dopa = L-3,4-dihydroxyphenylalanine. DDC = dopa decarboxylase. TAT = tyrosine aminotransferase. GOT1 = glutamate oxaloacetate transaminase 1. GOT2 = glutamate oxaloacetate transaminase 2. BH₄ = tetrahydrobiopterin. BH₂ = dihydrobiopterin.

### Serum metabolomic analysis from PKU patients

Given the importance of L-tyrosine for neurotransmitter synthesis and also the reported variation in neurological outcomes of well-treated PKU patients ^13^, we further investigated the role of L-tyrosine in PKU. Therefore, we re-analysed data from a recent targeted metabolomic study on 27 patients (ages 1.6–48.6 years old, **Figure 2A**) with confirmed PKU and 32 healthy controls (ages 19–60 years old) ^13^. As expected, the measured L-phenylalanine levels of the PKU patients were higher in patients than in controls (**Figure 2B**). However, the measured L-tyrosine levels were also significantly higher in PKU patients than in healthy controls (**Figure 2B**). This result is consistent with the original paper but contradicts our earlier predictions and other reports ^3^. All PKU patients have received a diet consisting of natural protein and L-phenylalanine-free medical foods containing amino acid mixtures as well as L-tyrosine supplementation ^13^, which may explain the high L-tyrosine levels. No relationship could be found between estimated L-phenylalanine intake and measured serum L-phenylalanine or L-tyrosine levels (**Figure 2B**, C), which was also consistent with another study ^35^. Moreover, while the serum L-phenylalanine levels in healthy individuals correlated with the L-tyrosine levels (R^2^ = 0.57), such correlation was not observed for the PKU patients (R^2^ = 3e^-3^). If no external factors contributed to the L-tyrosine levels, one would have expected a negative correlation between L-phenylalanine and L-tyrosine. However, this lack of correlation could be due to the medical diet, as suggested by Moritz et al. Another contributing factor could be the gut microbiome, which we decided to further investigate using our infant-WBMs.

**Figure 2:**
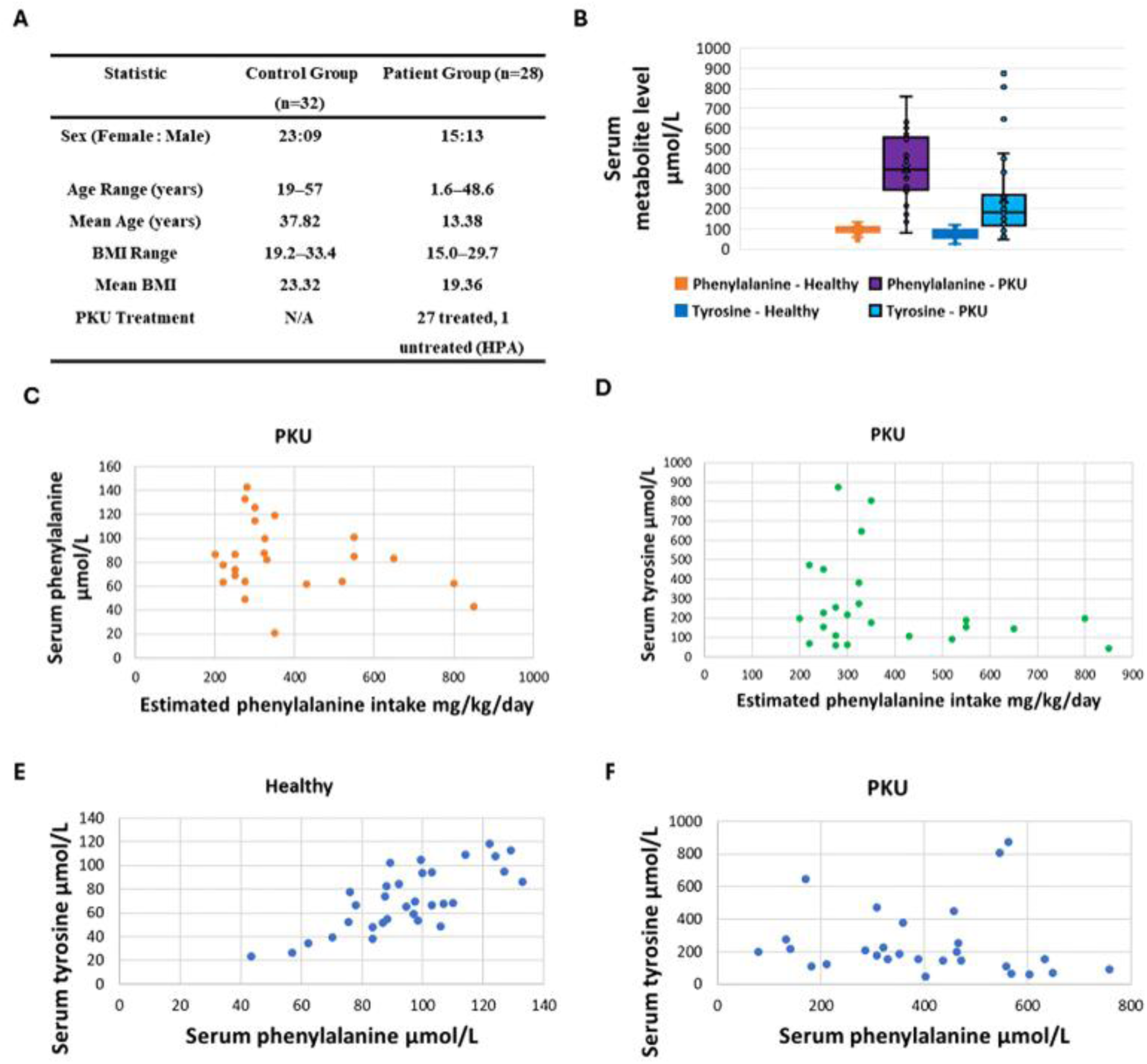
Serum metabolomics and L-phenylalanine intake analysis in PKU patients and healthy controls. (A) Summary table comparing control and patient groups (n=32 and n=28, respectively) based on sex distribution, age range, mean age, BMI range, mean BMI, and treatment status. (B) Serum L-phenylalanine and L-tyrosine levels in PKU patients versus healthy controls. (C) Relationship between estimated L-phenylalanine intake and serum L-phenylalanine levels in PKU patients. (D) Relationship between estimated L-phenylalanine intake and serum L-tyrosine levels in PKU patients. (E) Correlation between serum L-phenylalanine and L-tyrosine levels in healthy controls. (F) Correlation between serum L-phenylalanine and tyrosine levels in PKU patients. HPA = Hyperphenylalaninemia.

### Impact of microbiome on *in silico* biomarker levels

No matching gut microbiome data was available for the PKU patients from the metabolomics study. Hence, to address the question of whether the gut microbiome could affect L-tyrosine blood levels in PKU, we obtained microbial community models from the APOLLO resource ^33^ of 48 female infants aged from 2 to 13 days old (mean = 3.9 days, standard deviation (±) 2.86 days) at the time of sample selection, with 16 being from infants at 2 days. None of the selected infants had received antibiotics and the samples were from 41 vaginally and seven C-section-delivered infants (**Table S2**). We included both breast and formula-feed infants (**Table S2**). The microbial community models have been built based on the APOLLO microbial reconstructions and published metagenomic data from healthy female infants ^36^ (Methods). Overall, the microbial community models contained on average 118,350 reactions and 109,060 metabolites (**Table 1**).

**Table 1.** Genome-scale reconstructions scope. This table summarises the reaction and metabolite content of the computational models used in this study. std = standard deviation. NA = not applicable.

| Reconstruction type | # reactions (std) | # metabolites (std) | # microbial species |
| --- | --- | --- | --- |
| Germ-free female infant-WBM (Age 2 days) | 85,662 | 60,436 | 0 |
| Microbial community models (n=48) | 118,350 ($\pm 40,385$) | 109,060 ($\pm 39,035$) | 100.8 ($\pm 42.5$) |
| Microbiome female infant-WBMs (n=48) | 201,990 ($\pm 40,385$) | 168,270 ($\pm 39,035$) | 100.8 ($\pm 42.5$) |

#### Compositional diversity of the 48 infant gut microbiomes

We investigated the microbial composition of the 48 gut microbiomes of female infants underlying the microbial community models (**Figure 3**). The 48 microbiomes consisted of 789 unique microbes from seven phyla (**Table S3**). Each sample contained on average 100.8 ± 42.5 species (with a range from 49 to 232 species). The Pielou’s evenness, a measure of microbial diversity within a sample, ranged from 0.41 and 0.9 with an average of 0.78 ± 0.08, illustrating the variety in the composition of the microbiomes (**Table S4**). On average, most bacterial species belonged to Firmicutes (41.6 ± 24.6 %), followed by Bacteroidetes (28.2 ± 20.6 %) and Actinobacteria (17.5 ± 12.1 %), with smaller proportions of Proteobacteria and others (**Figure 3**). Most phyla were present in 30 or more samples, except for Fusobacteria and Verrucomicrobia, which were only present in one and two samples, respectively (**Table S5**). The differences between samples were even more pronounced on genus level, with only four out of 62 taxa being shared between at least 40/48 samples (i.e., Bacteroides, Bifidobacterium, Staphylococcus, and Streptococcus), but at highly different abundances. Most genera were only present in a few samples, with 27 of them in less than ten samples (**Table S6**). In conclusion, the microbial community samples and their microbial community metabolic models differed substantially in their microbial composition. Thus, these models were ideally suited to investigate whether the microbial metabolic activity could affect *in silico* the reliability of our predicted biomarkers in the infant-WBMs.

**Figure 3:**
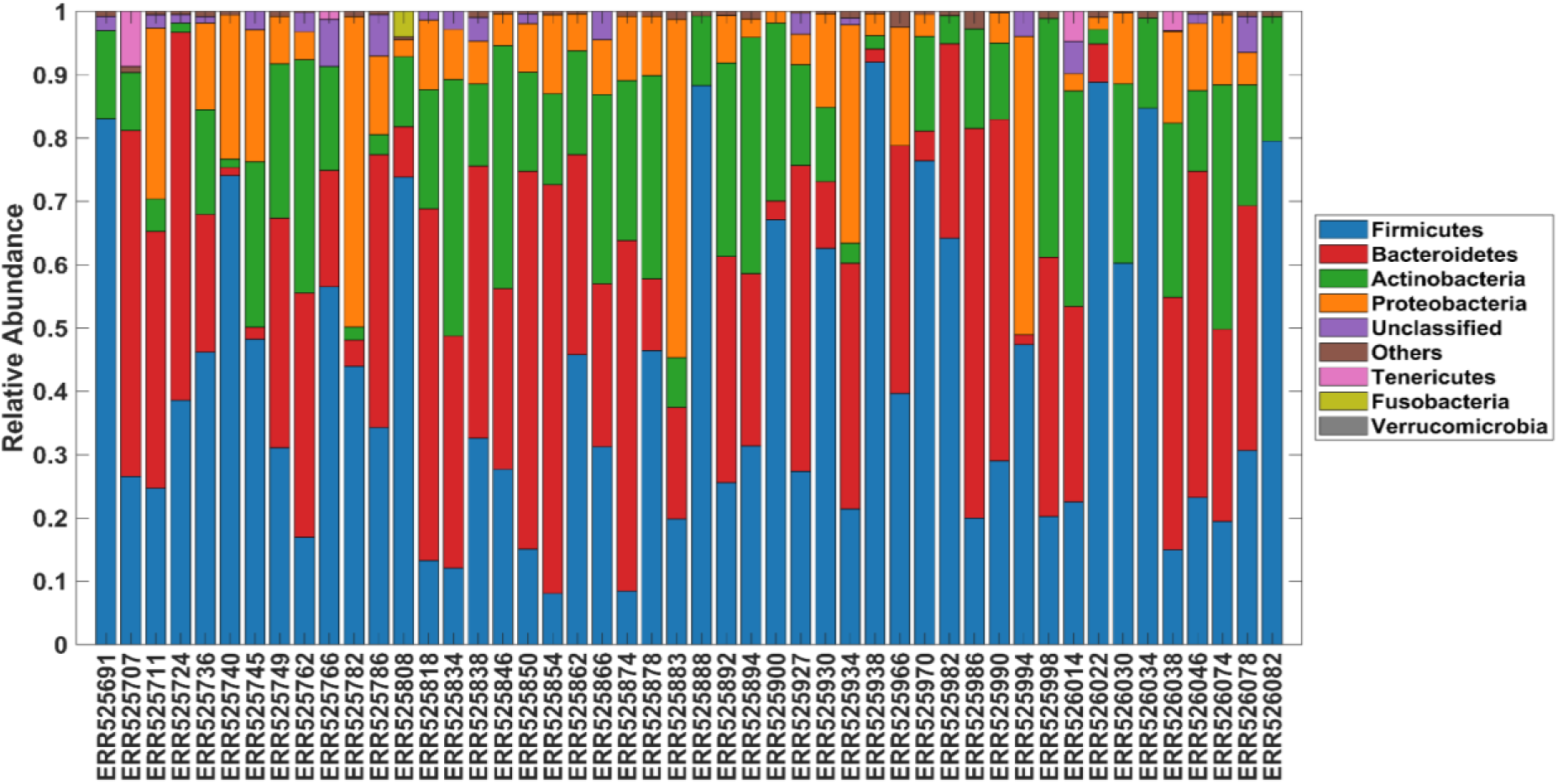
Relative abundance of microbial phyla across 48 gut metagenomics samples of healthy human female infants^33,36^.

#### Generation of microbiome infant-WBMs

Next, to create microbiome-associated WBMs, we placed each of the 48 microbial community models into the large intestinal lumen of the female infant-WBM (Methods) resulting in 48 microbiome infant-WBMs (**Table 1**). Since these microbiome infant-WBMs only differed in their microbial community composition (and the corresponding relative strain abundances), any predicted changes in the metabolic outputs were solely attributable to alterations in microbial composition. As above, the PKU models were created by setting the PAH-associated reaction bounds to zero, resulting in a total of 96 microbiome infant-WBMs, i.e., 48 healthy and 48 corresponding PKU models.

#### Microbial metabolism alters predicted biomarkers

Next, we asked whether the identified biomarkers in L-phenylalanine and L-tyrosine metabolism using the germ-free infant-WBM would remain reliable despite possible variation in microbial metabolic activity due to differences in microbiome composition. Therefore, we recomputed the maximal possible flux value for all candidate biomarkers using the 48 healthy microbiome infant-WBMs. We found that in all cases, the presence of the microbiome models led to substantially higher flux values (**Table 2**). We then repeated the simulations using the 48 PKU microbiome infant-WBMs and found that the maximal possible fluxes for the established PKU biomarkers L-phenylalanine, hydroxyphenylacetic acid, and phenylpyruvic acid remained significantly increased (p-value < 1e^-21^) (**Figure 4**, **Table 2**). In contrast, the maximal possible fluxes for L-tyrosine and its derivatives were not statistically significant anymore, except for L-tyrosine (p=4e^-3^, **Figure 4**). These results demonstrate that microbial metabolism could alter the reliability of blood metabolites as biomarkers due to host-microbiome co-metabolism.

**Figure 4.**
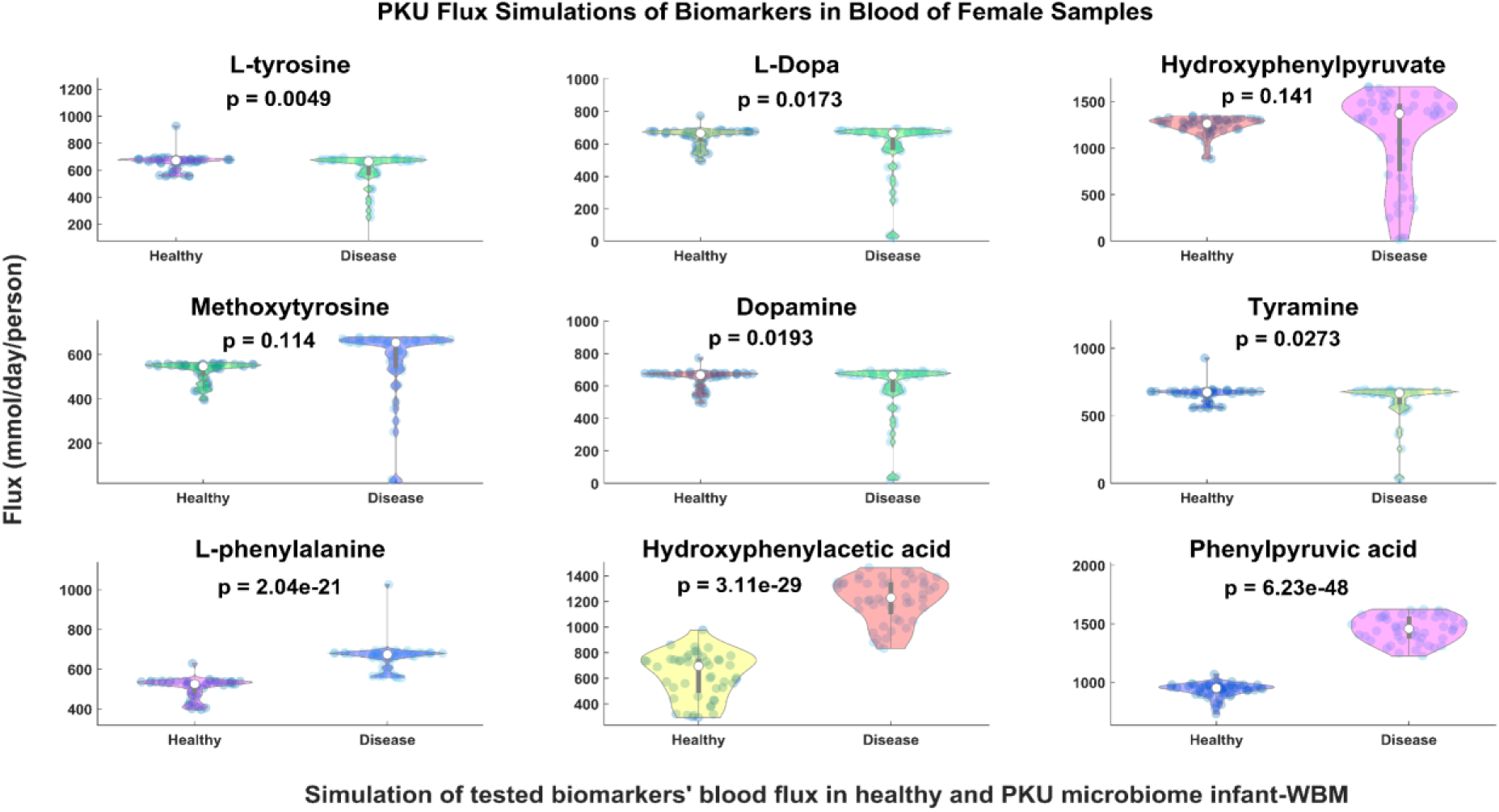
Predicted maximal possible blood fluxes for the metabolites along the L-phenylalanine and L-tyrosine metabolism using the 48 healthy and 48 PKU (“Disease”) microbiome infant-WBMs.

**Table 2.** Predicted biomarker fluxes under healthy and PKU conditions in germ-free and host-microbiome infant-WBMs. The predicted fluxes of all tested biomarkers are shown, highlighting the differences between germ-free and host-microbiome simulations. Flux units are given in mmol/infant/day. Std = standard deviation. VMH = Virtual metabolic human database ^37^.

| Biomarker | VMH ID | Healthy germ-free flux | PKU germ-free flux | Average healthy host-microbiome flux (std) | Average PKU host-microbiome flux (std) | p-value |
| --- | --- | --- | --- | --- | --- | --- |
| L-tyrosine | tyr_L | 8.06 | 0.19 | 656.32 (± 60.29) | 578.23 (± 177.8) | 0.0049 |
| L-Dopa | L-dopa | 8.06 | 0.19 | 643.08 (±56.77) | 577.83 (± 177.6) | 0.0173 |
| 3,4-hydroxyphenylpyruvate | 34hpp | 8.06 | 0.19 | 1,236.83 (± 103.1) | 1,127.67 (± 499.5) | 0.141 |
| Methoxytyrosine | CE2176 | 8.06 | 0.19 | 523.4 (± 46.5) | 564.53 (± 172.6) | 0.114 |
| Dopamine | dopa | 8.06 | 3.47 | 642.52 (± 56.79) | 578.78 (± 176.6) | 0.0193 |
| Tyramine | tym | 8.06 | 3.47 | 656.3 (± 60.18) | 604.17 (± 149.4) | 0.0273 |
| L-phenylalanine | phe_L | 1.97 | 7.87 | 505.64 (± 50.42) | 659.64 (± 70.05) | 2.04e-21 |
| Hydroxyphenylacetic acid | 2hyoxplac | 1.97 | 7.87 | 621.9 (± 179.8) | 1,211.97 (± 173.7) | 3.11e-29 |
| Phenylpyruvic acid | phpyr | 1.97 | 7.87 | 934.86 (± 61.03) | 1,456.32 (± 111.5) | 6.23e-48 |

### Impact of microbial metabolism on predicted flux values for L-tyrosine and its derivatives

Since the predicted L-tyrosine flux values varied substantially between the PKU microbiome infant-WBMs (**Figure 4**), we further analysed our predictions to establish whether L-tyrosine could serve as a reliable biomarker. In 37 out of 48 PKU microbiome infant-WBMs, the L-tyrosine fluxes were unchanged when compared to their respective healthy counterparts (**Table S7**). This result means that microbially produced L-tyrosine contributed to the host’s L-tyrosine availability in these 37 models. In contrast, in 11 PKU microbiome infant-WBMs, the maximal possible L-tyrosine flux was drastically reduced. For example, in one of the individuals (ERR525982), the maximal possible L-tyrosine flux was only 2.8% of that achieved in its healthy microbiome infant-WBM counterpart (**Figure 4**, **Table S7**). In these 11 models, gut microbial metabolism failed to compensate for PAH-deficiency, likely due to the absence or insufficient abundance of certain microbial taxa required for microbial L-tyrosine biosynthesis. Four of these 11 (36%) infants were born through Caesarean section (CS). The other three infants born through CS showed lower L-tyrosine production capabilities in the healthy case compared to the vaginally born infant-WBMs, but no further reduction in predicted blood L-tyrosine flux in the PKU simulations was observed (**Table S1**). In contrast, no correlation with sampling day after delivery or feeding regime (breast vs formula-fed) could be observed. Similarly, no difference in species richness and Pielou’s evenness was found. However, we noted that seven of the eleven microbiomes had a summed relative abundance of >70% of Firmicutes (**Figure 3**, **Table S3**). Taken together, these results demonstrate that microbial metabolism could lead to interindividual variations, which may affect the suitability and reliability of L-tyrosine, and its derivatives, as PKU biomarkers, and suggest a subpopulation, for which such biomarker may be suitable.

#### Microbes improving in silico L-tyrosine production

To identify key contributing L-tyrosine-producing microbes in the 11 L-tyrosine-decreased PKU microbiome infant-WBMs, we used a feature of linear programming called shadow price. Briefly, each time a linear program is solved, i.e., in our case the maximisation of blood L-tyrosine flux in a microbiome infant-WBM, the shadow price is also calculated for each metabolite in the model. The shadow price represents how much the maximal flux value for objective function (here: blood L-tyrosine reaction) would increase if one were to increase the flux through this metabolite. As each microbe’s biomass was effectively treated as a metabolite in the infant-WBMs (represented through its biomass and the associated biomass reaction), we obtained the shadow price for each microbe and its effect on the maximal possible blood L-tyrosine flux value (Methods). For our analysis, we only considered microbes that were present in more than ten percent of the 48 microbiome infant-WBMs, i.e., 252 out of 789 (32%) microbes and performed a Z-score analysis on the shadow price values of these 252 microbes across the 48 PKU microbiome infant-WBMs. Microbes with a Z-score less than −1 were classified as having *strongly negative shadow prices* (SNSPs). This threshold was chosen to highlight microbes with the strongest potential to increase the flux through the blood L-tyrosine reaction. Among all tested microbes, 16 out of 252 (6.3%) exhibited SNSPs (**Figure 5**). We classified these 16 microbes as critical microbes for tyrosine synthesis. From the top 10 microbes (Table 3), two were excluded, being *Bacteroides fragilis* 3_1_12 and *Rothia* SRR5963194_bin_1, as they had no SNSPs in some of 11 L-tyrosine-decreased PKU microbiome infant-WBMs (**Table 3**). Note that *Bifidobacterium dentium* ATCC 27678 remained in the target list, as it was absent from all L-tyrosine-decreased PKU microbiome infant-WBMs. *Propionibacterium sp.* 20298_3_1 had a higher abundance in the L-tyrosine-decreased PKU microbiome infant-WBMs compared to PKU microbiome infant-WBMs with normal L-tyrosine flux (**Figure 6**). Therefore, we also excluded this microbe from the list of potential critical microbes for L-tyrosine synthesis. From the remaining seven microbes, three microbes were unclassified, belonging to the *Bifidobacterium* and the *Escherichia* genera, while the remaining three named species belonged to the *Bifidobacterium* genus (**Table 3**). Additionally, we determined whether there were any microbes with strongly positive shadow prices, but could not find such microbes in the PKU microbiome infant-WBMs. Taken together, we identified three unclassified and four classified species that may be critical for L-tyrosine metabolism.

**Figure 5.**
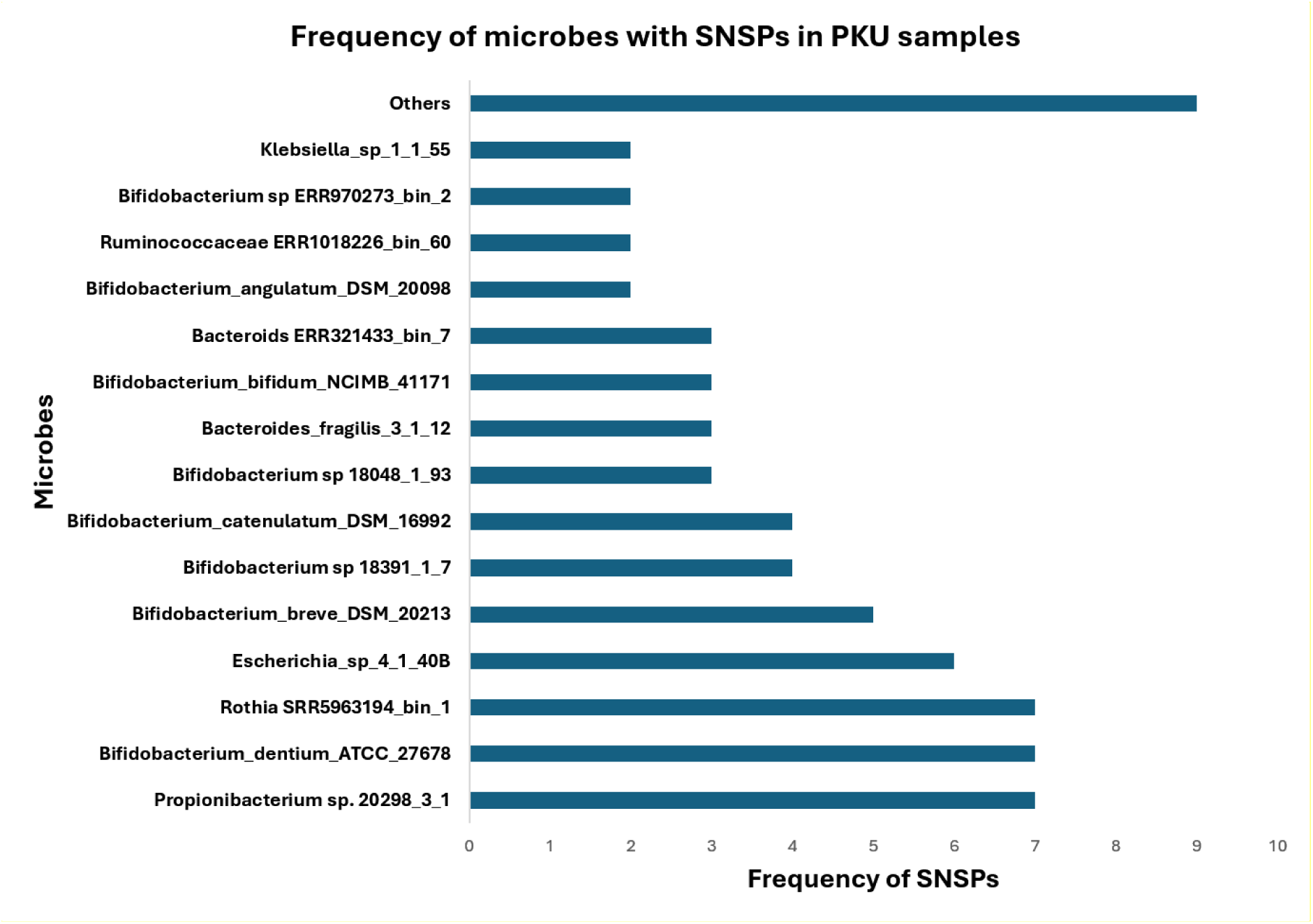
Frequency of 16 microbes with strongly negative shadow prices (SNSPs) across the 48 PKU microbiome infant-WBMs with respect to the maximal possible blood L-tyrosine flux. These microbes were classified as *critical microbes for L-tyrosine synthesis*.

**Figure 6.**
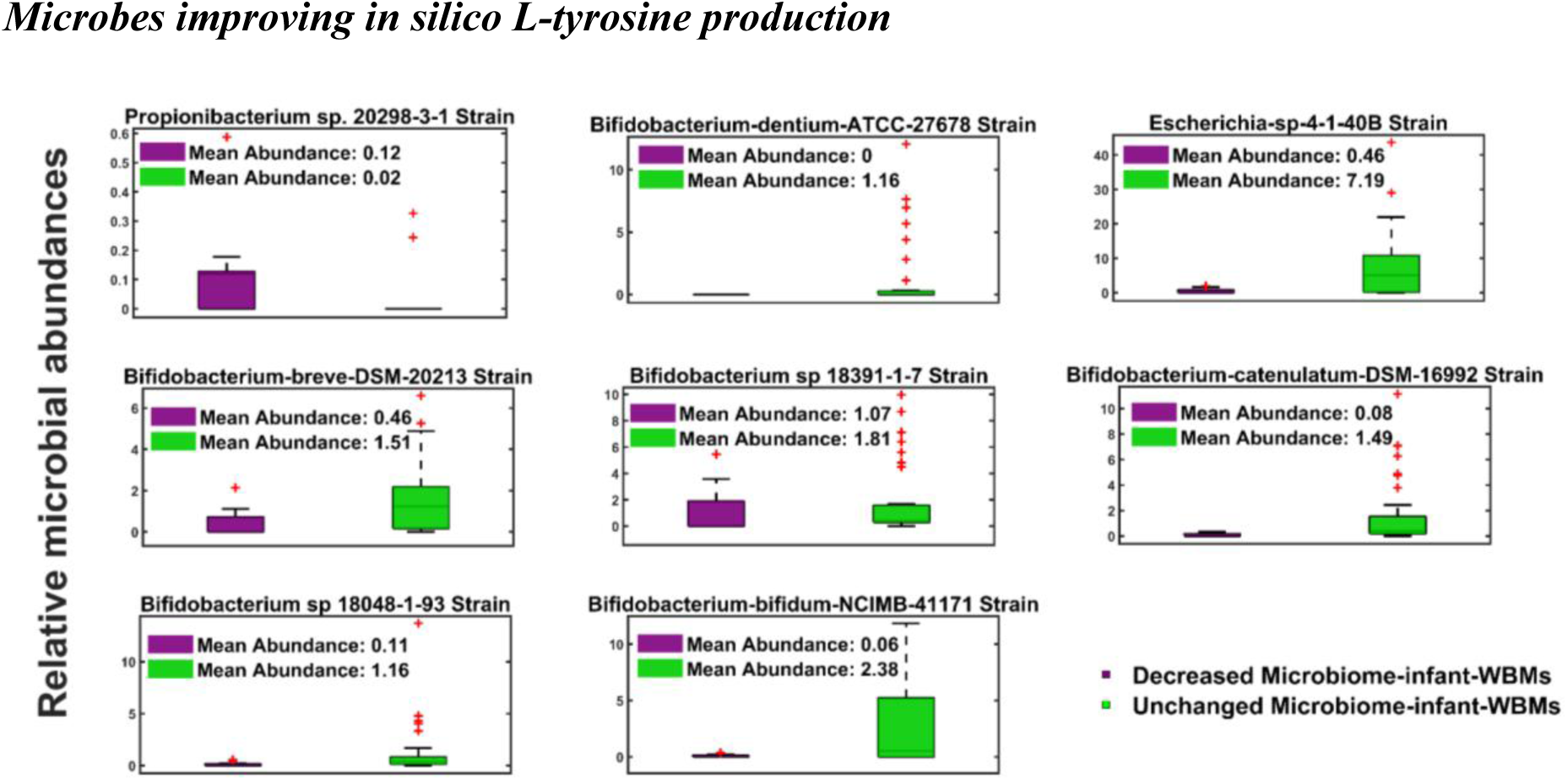
**Comparison of the relative abundances of the key contributing L-tyrosine-producing microbes** in the 11 microbiome infant-WBMs with predicted decreased L-tyrosine blood flux values and 37 microbiome infant-WBMs with predicted unchanged L-tyrosine blood flux values under PKU simulations.

**Table 3.**
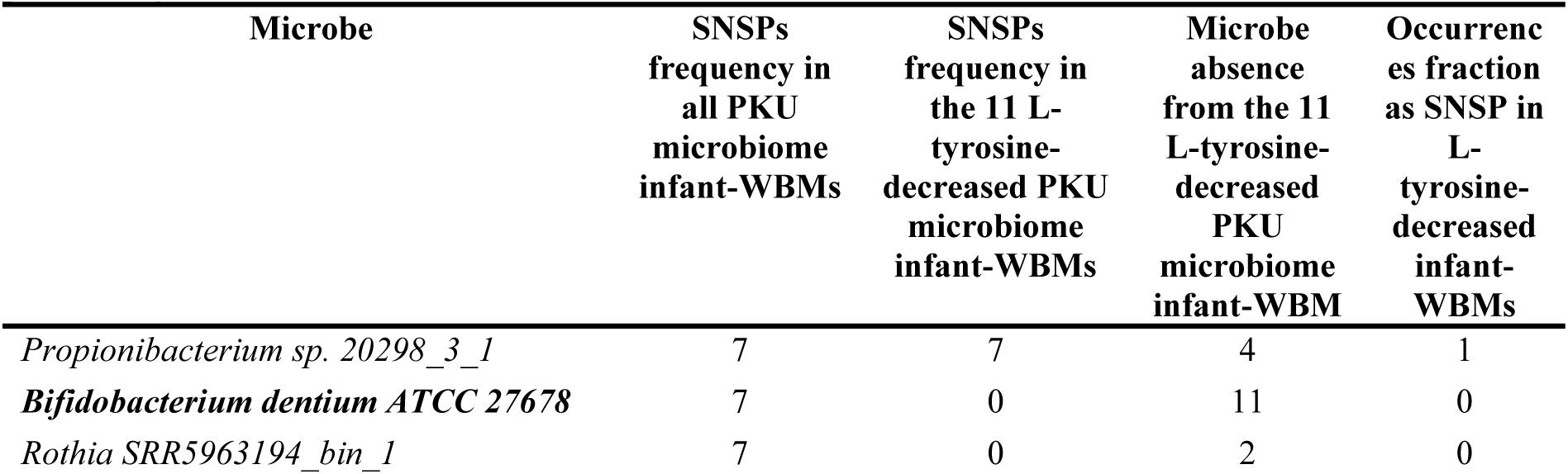
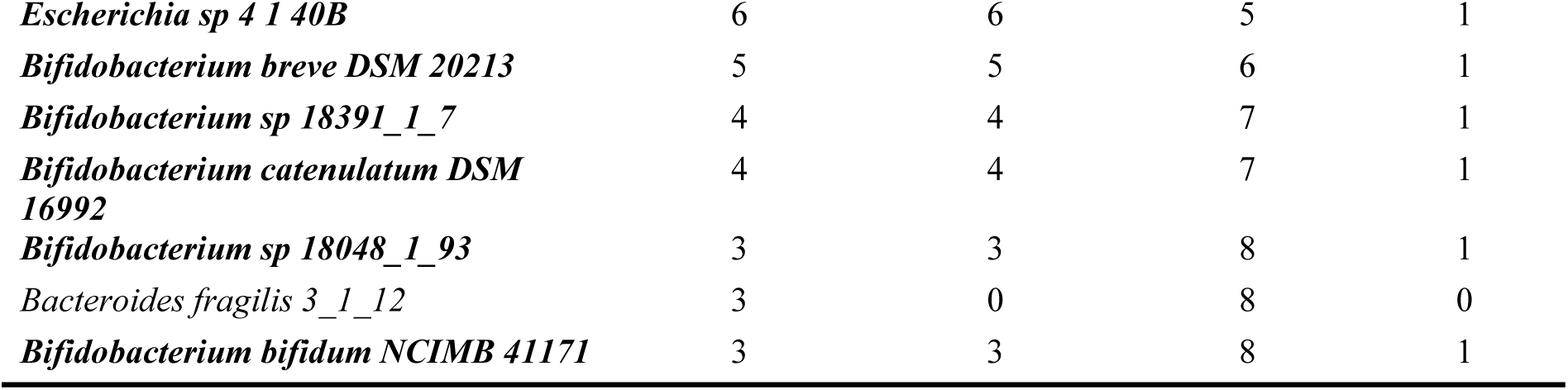
Strongly negative shadow price (SNSPs) and abundance patterns of microbes in the microbiome infant-WBMs when maximising for L-tyrosine flux in the blood compartment. This table summarises the occurrence of SNSP for microbial strains across 48 microbiome infant-WBMs. The second column shows the total number of SNSP occurrences across 48 PKU microbiome infant-WBMs, while the third column highlights the occurrences exclusive to the 11 L-tyrosine-decreased PKU microbiome infant-WBMs. The fourth column reports the number of PKU microbiome infant-WBMs, where the microbe was absent, and the fifth column provides the fraction of L-tyrosine-decreased PKU microbiome infant-WBMs where the microbe was identified as microbes exhibiting SNSP. Key contributing L-tyrosine-producing microbes are bold.

#### Microbial L-tyrosine synthesis route explains the predicted L-tyrosine reduction

As our results demonstrated that 77% of the infant microbiome could compensate for the defective PAH gene in the WBM, we further investigated how the microbiome models achieved this, and which reactions (or pathways) may be absent or less abundant in the identified strains. Overall, in bacteria, the biosynthesis of L-tyrosine can occur via three pathways (**Figure 7**). The first two pathways begin with the conversion of chorismate, which originates from the Shikimate pathway, into prephenate by the chorismate dismutase (VMH ID: CHORM) (**Figure 7A**). In the hydroxyphenylpyruvate pathway, prephenate is then converted into 4-hydroxy-phenylpyruvate by the prephenate dehydrogenase (VMH IDs: PPND, PPND2), which is further metabolised by the L-tyrosine transaminase (VMH ID: TYRTA) into L-tyrosine. Alternatively, via the pretyrosine pathway, prephenate can be converted into pretyrosine (arogenate) by the L-arogenate aminotransferase (VMH IDs: LAGAT, LARGNAT). Arogenate is then converted into L-tyrosine by the arogenate dehydrogenase (VMH IDs: L_AROND, L_ARONR) ^38^. In the third pathway, L-phenylalanine is converted into L-tyrosine by the phenylalanine hydroxylase (VMH ID: PHETHPTOX2) ^39^ (**Figure 7A**). Relative reaction abundances were determined in each microbiome infant-WBM by adding the relative abundance of each microbe in the microbial community model, which carries the reaction. The summed relative abundance of the reactions along the hydroxyphenylpyruvate pathway did not differ between PKU models with normal L-tyrosine blood flux and those with reduced L-tyrosine blood flux (**Figure 7B**). In contrast, the summed relative abundances of the reactions involved in the pretyrosine pathway were statistically significantly reduced in the 11 L-tyrosine-decreased PKU microbiome infant-WBMs (**Figure 7B**). Notably, the L-arogenate aminotransferase reaction (VMH ID: LAGAT) was completely absent in the 11 L-tyrosine-decreased PKU microbiome infant-WBMs (**Figure 7B**). The third pathway (PHETHPTOX2) was absent from all microbial community models, which was not surprising as only 146 microbial metabolic reconstructions of the 7,321 one contained in the AGORA2 resource contained these reactions. Moreover, most (137/146) of these reconstructions belonged to proteobacteria. As a next step, we confirmed that all of the identified key contributing L-tyrosine-producing microbes contained the pre-tyrosine pathway. Moreover, in 37 PKU microbiome infant-WBMs with normal L-tyrosine, these strains produced more than 50 percent of the total L-tyrosine flux (**Table S8**), mainly through the pretyrosine pathway. In contrast, in the 11 L-tyrosine-decreased PKU microbiome infant-WBMs, the flux through this pathway was lower. These results confirm that lower potential in microbial L-tyrosine compensation could be attributed to a lower abundance of strains encoding the pretyrosine pathway.

**Figure 7.**
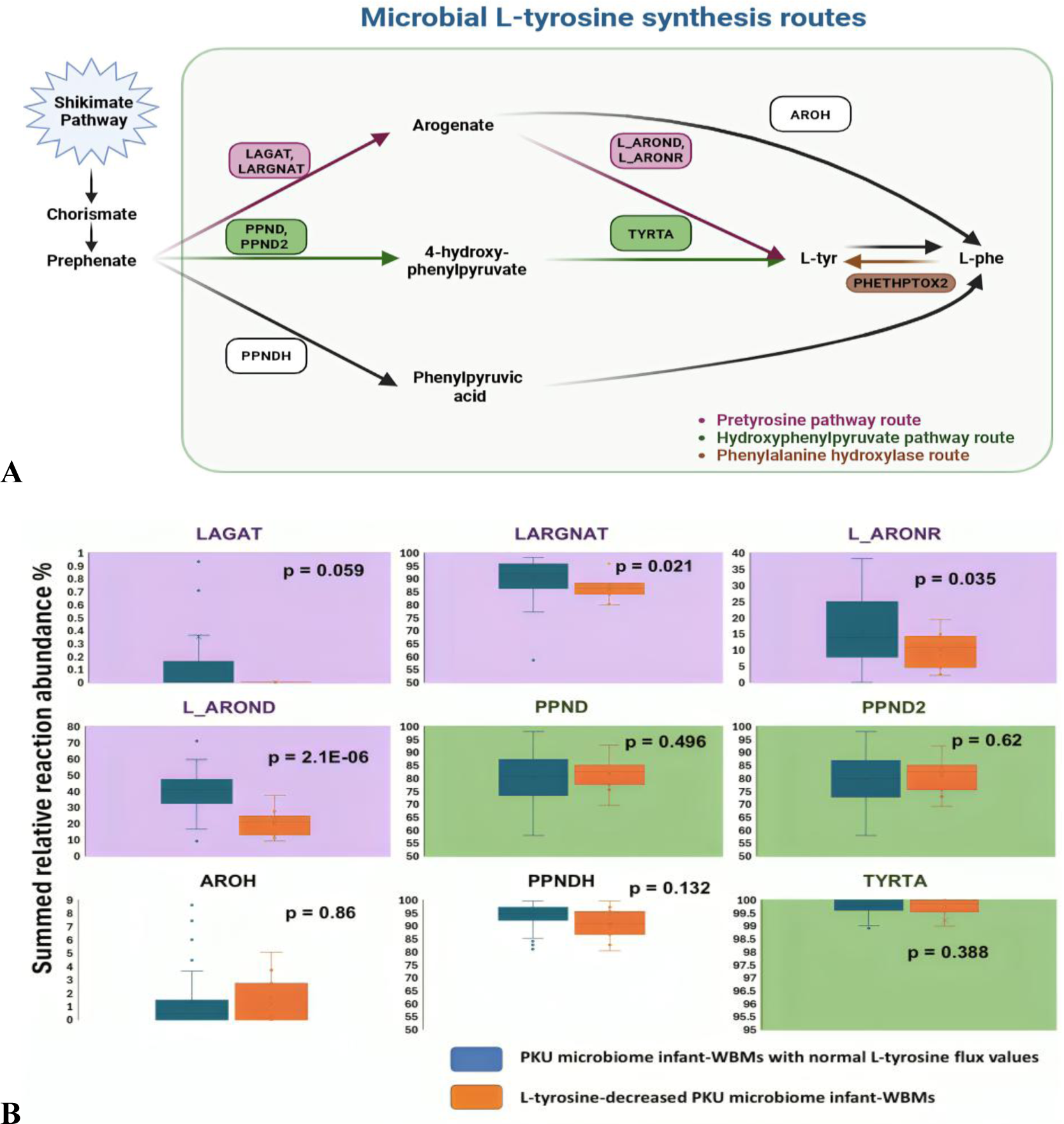
Microbial pathways for L-tyrosine biosynthesis. **A.** Three microbial routes to L-tyrosine are shown: (1) the hydroxyphenylpyruvate pathway via prephenate dehydrogenase (PPND/PPND2) and L-tyrosine transaminase (TYRTA), (2) the pretyrosine pathway via L-arogenate aminotransferase (LAGAT/LARGNAT) and arogenate dehydrogenase (L_AROND/L_ARONR), and (3) the phenylalanine hydroxylase route (PHETHPTOX2). **B.** Relative microbial reaction abundances in the 37 PKU microbiome infant-WBMs with normal L-tyrosine flux values and the 11 L-tyrosine-decreased PKU microbiome infant-WBMs.

### Microbes that affect host phenylalanine flux

Finally, we investigated whether any gut microbes could counteract the protein-reduced diet and phenylalanine-free medical food supplementation by altering L-phenylalanine bioavailability for the host. In our earlier results (**Figure 1**, **Figure 4**), we predicted an average increase of L-phenylalanine flux of 154 ±36.8 mmol/day/infant in the presence of the microbiome models in the infant-WBMs compared to the germ-free infant-WBMs. Hence, we repeated the shadow price analysis, this time looking for microbes with substantial positive shadow prices, which would correspond to an ability to reduce blood L-phenylalanine levels, e.g., by consuming more of the dietary L-phenylalanine or microbially produced L-phenyalanine within the community. However, we could not identify such microbes. In contrast, we found several microbes, which had SNSPs, suggesting that these microbes could increase L-phenylalanine production substantially if present at higher relative abundance. To identify key L-phenylalanine contributors, we repeated the Z-score analysis and identified all microbes with a Z-scores less than −1. Among the four identified microbes with substantial contributions to L-phenylalanine production (**Table 4**), one was an unspecified microbial strain (*Rothia* SRR5963194_bin_1). The remaining key L-phenylalanine contributors were *Bifidobacterium dentium ATCC 27678, Bacteroides fragilis 3_1_12*, and *Bacteroides caccae* ATCC 43185. Taken together, we could not identify any microbes that could individually substantially reduce the blood L-phenylalanine flux. However, we identified four microbes that could further increase the L-phenylalanine flux, thereby potentially worsening the phenotype of the PKU patients and counteracting dietary therapy.

**Table 4.** SNSPs patterns of microbes in the microbiome infant-WBMs when maximising for L-phenylalanine production in blood. This table summarises the occurrence of SNSP for microbial strains across all 48 PKU microbiome infant-WBMs. The second column shows the total number of SNSP occurrences across all PKU microbiome infant-WBMs. The third column reports the number of microbiome infant-WBMs with predicted increased L-phenylalanine blood flux value, where the microbe was absent, and the third column provides the fraction of phenylalanine-increased infant-WBMs where the microbe was identified as exhibiting SNSP.

| Microbes | SNSPs frequency in all PKU simulated microbiome infantWBMs | Number of L-phenylalanine-increased microbiome infant-WBMs, where microbe was absent | Occurrence fraction as SNSP in L-phenylalanine-increased microbiome infant-WBMs |
| --- | --- | --- | --- |
| *Rothia* SRR5963194_bin_1 | 8 | 17 | 0.52 |
| *Bifidobacterium dentium* ATCC 27678 | 7 | 30 | 0.71 |
| *Bacteroides fragilis* 3_1_12 | 4 | 16 | 0.42 |
| *Bacteroides caccae* ATCC 43185 | 1 | 19 | 0.42 |

## Discussion

In this study, we aimed at identifying reliable biomarkers for PKU using whole-body metabolic modelling. To achieve this, we used a published germ-free infant whole-body metabolic model (infant-WBM) to first predict biomarkers in L-phenylalanine and in L-tyrosine metabolism. While L-phenylalanine and phenylpyruvic acid were known disease biomarkers ^34^, hydroxyphenylacetic acid and L-tyrosine, and its derivatives, are potentially novel biomarkers. However, changes in blood levels of L-tyrosine have been reported to be lower or unchanged in PKU patients, which could be either due to its dietary supplementations and/or gut microbial L-tyrosine metabolism. To further investigate the natural variation of L-tyrosine as a function of the gut microbiome, we used published infant gut microbiome metabolic models ^33^ and integrated them with the infant-WBMs ^31^. We found that indeed the microbiome could contribute to host L-tyrosine metabolism in most, but not all cases. Consequently, we identified candidate microbes associated with low L-tyrosine production potentials, whose absence or lower abundance would result in lower blood L-tyrosine levels. As many of these microbes belonged to the *Bifidobacterium* genus, they may be used for a probiotic treatment of PKU to supplement L-tyrosine to achieve healthy levels. Taken together, we demonstrate how microbiome infant-WBMs and metabolic modelling can be used to predict candidate novel biomarkers, while considering interindividual variations, and point towards candidate probiotic agents.

To-date, multiple metabolic modelling studies have predicted known and novel biomarkers for IMDs ^40^. In line with those studies, we predicted two known and potentially seven novel biomarker metabolites (**Figure 1**), among a set of 233 metabolites (**Table S1)**, all of which either belonged to the L-phenylalanine or the L-tyrosine metabolism. Comparison of our predicted metabolites with metabolomic results ^13^, and in accordance with other reports ^3^, revealed that L-tyrosine and its derivatives are not altered in all PKU patients (**Figure 2**). These results raise the questions on how much the metabolic modelling approach could be used to identify reliable biomarker metabolites. This shortcoming exists because available human models did not account for interindividual variability caused by genomic variations, different life-style conditions, or the gut microbiome. In our study, we overcame the shortcoming by accounting for interindividual variability due to difference in gut microbiome composition.

The adult WBMs have already been expanded with personalised microbiome models ^30,41,42^ to demonstrate their value in prediction of known and novel host-microbiome co-metabolism. In the present study, we extended the published infant-WBMs (day 2) with published infant gut microbiome models, which allowed us to assess the reliability of the predicted PKU biomarkers. As expected, the established biomarkers for PKU, L-phenylalanine and phenylpyruvate, remained statistically significant in the presence of the microbial community models in the infant-WBMs (**Figure 4**). This result demonstrates the reliability of these biomarkers but also underlines that the modelling approach could be used to assess the consequences of interindividual microbiome variations for biomarker reliability. In contrast to L-phenylalanine, the predicted candidate biomarkers along the L-tyrosine metabolism changed with the variation of the gut microbial composition and thus, were not reliable (**Figure 4**). In fact, only in a quarter of the microbiome-associated PKU infant-WBMs, the predicted L-tyrosine blood flux value was lower than in the healthy counterpart. Our analysis of differences in microbial composition of those microbiomes revealed an enrichment in Firmicutes and a depletion of Bifidobacterium (**Table 2**). Such infant gut microbial variation can result from, e.g., differences in milk composition ^43^ and delivery mode^44^. Accordingly, most infant microbiomes with increased Firmicutes occurred in C-section delivered infants, which were also enriched in the 11 L-tyrosine-decreased PKU microbiome infant-WBMs (**Table S8**). Similarly, reduced Bifidobacterium relative abundance has been reported for C-section delivered infants ^45^. We suggest, based on our simulation results, that this infant population might be at a higher risk of decreased L-tyrosine blood levels and for which L-tyrosine may be a suitable secondary biomarker for PKU. Additionally, once a PKU diagnosis is confirmed, these infants may need a higher L-tyrosine supplementation than other PKU infants.

On a pathway level, we could establish that those L-tyrosine-decreased PKU microbiome infant-WBMs had missing or lower abundances of the reactions along the pre-tyrosine pathway (**Figure 7**). This result provides a mechanistic explanation for predicted reduced L-tyrosine levels based on microbial metabolic activity and demonstrates how microbial metabolism can complement human metabolism, as this pathway is absent in humans. In plants, the pre-tyrosine pathway is the predominant pathway for L-phenylalanine synthesis, due to the high catalytic efficiency of the prephenate aminotransferase ^46^. While this pathway has been studied in plants and cyanobacteria ^46^, its role in microbes remains understudied.

In particular, we identified six Bifidobacterium strains to be responsible for the predicted lower tyrosine blood fluxes (**Figure 6**). Beyond their role in amino acid metabolism, Bifidobacterium species offer additional health benefits. For instance*, B. bifidum* is known to support immune function by enhancing the body’s defence mechanisms and inhibiting the growth of harmful bacteria ^47^. Additionally, *B. catenulatum* contributes to maintaining gut health by producing short-chain fatty acids, which lower intestinal pH and inhibit the growth of pathogenic bacteria ^48^. These benefits underscore the multifaceted advantages of Bifidobacterium strains in promoting overall health. However, our analysis also revealed that *B. dentium*, despite its potential for tyrosine production, could also contribute to increased L-phenylalanine levels, which may render it unsuitable as a therapeutic strain (**Table 3**). This finding indicates the importance of carefully evaluating the dual effects of microbial strains when considering their applications in PKU management, as it has been suggested in the literature ^49^. Nonetheless, the multifaceted advantages of Bifidobacterium strains, including their proven efficacy in enhancing gut health, immune support, and metabolic balance, alongside their potential to address L-tyrosine deficiencies, position them as critical targets for microbiome-based therapies.

In addition to Bifidobacterium, our simulations identified *Escherichia sp. 4_1_40B* as another microbial strain with the potential to decrease L-tyrosine deficiencies in PKU patients (**Table 2**). While this strain has not been extensively studied in the context of PKU, its metabolic capabilities and contributions to L-tyrosine production in our models support further investigation. Similarly, two unspecified microbial strains, *18391_1_7* and *18048_1_93*, both belonging to the Bifidobacterium genus, emerged as significant contributors to L-tyrosine production in the PKU infant-WBMs. These findings emphasise the importance of advancing research on the gut microbiome in newborns, particularly focusing on unclassified or poorly characterised species. The identification and functional characterisation of these strains could reveal novel therapeutic targets and deepen our understanding of how the gut microbiome influences inherited metabolic diseases, such as PKU.

Diurnal variations of L-phenylalanine and L-tyrosine blood concentrations in PKU patients has also been reported ^50–52^. While at least the L-tyrosine blood levels could be shown to increase with meal intake, which has been explained with high tyrosine supplementation ^50^. Interestingly, diurnal variations of up to 20% of the relative abundances of gut microbes have been reported ^53^, which could lead to changed composition of the entire community during a day ^54^. Again, dietary input has been suggested as one contributing factor ^53,54^, as well as hormone levels and activity levels of the human host. Moreover, at least in mice it has been shown that these intra-day variations in microbial composition could be connected to the host’s clock ^55^. Simulating such intra-day variations in diet uptake and microbial composition would be a valuable and insightful extension to our study. Such diurnal simulations could be achieved by combining COBRA modelling approach with kinetic modelling ^56–58^.

By using probiotics, prebiotics, or other microbiota-targeted interventions, researchers aim to lower L-phenylalanine levels and reduce the dietary restrictions imposed on PKU patients ^20^. For instance, it has been shown that genetically engineered *E. coli* Nissle could reduce the L-

phenylalanine blood levels in PKU mice ^59,60^. However, to our knowledge, using probiotics to manage L-tyrosine levels in PKU patients has not yet been reported. Probiotics could also aim at reducing L-phenylalanine producing microbes through competition or removal microbial produced L-phenylalanine. Similarly, probiotics could be used to eliminate microbes that compete for L-tyrosine or degrade L-tyrosine and thus lead to lower L-tyrosine blood levels. While we have not focused on identifying L-tyrosine reducing or competing microbes, our modelling approach would be suitable to also identify those ones. Overall, as research in this area progresses, microbiome-based therapies could complement traditional dietary management, improving metabolic control and overall quality of life for PKU patients. Integrating microbiome studies into biomarker research offers a promising avenue for advancing PKU management and improving patient outcomes and reducing deeper systemic imbalances ^61^.

In our study, we used microbiome data from healthy newborns in the absence of corresponding microbiome data from PKU newborns to simulate the consequences of the missing PAH gene on biomarker bioavailability and reliability. So far, most microbiome studies of PKU patients analysed stool samples from older-aged children ^62^ and adults ^63^. As the diet of these older patients is quite different than in newborns, reported microbial differences between PKU patients and healthy controls may not be transferable to newborns. It is not clear whether the gut microbiome composition of PKU newborns would differ significantly from healthy newborns, and whether any differences could then be attributed to the diseases. Hence, in the absence of PKU specific data, our approach seems reasonable.

Finally, the application of organ-resolved, infant-specific whole-body metabolic models integrated with personalised microbiome data offers a novel framework for the discovery and evaluation of novel, reliable biomarkers for any inherited metabolic diseases, as well as the development of possible targeted microbiome therapies.

## Material and method

### Computational models of infant whole-body metabolism (infant-WBMs)

We used the published female infant-WBM corresponding to a 2-day old infant from a resource of 360 infants ^31^, obtained from (https://doi.org/10.7910/DVN/SGQUEH). We used the pre-set constraints of the infant-WBM, if not specified differently. The female infant-WBM consisted of 1,724 unique genes (2,071 transcripts), 85,662 reactions, and 60,436 metabolites 30 organs and cell types and 13 biofluid compartments. The predefined diet consisted of human breast milk ^31^. The lower and upper bounds on the whole-body maintenance reaction were set to 1. Note that these infant-WBM were germ-free as no microbiome models had been associated with them.

### Metabolomic data

A published targeted metabolomic dataset of 27 PKU patients, one untreated patient with hyperphenylalanaemia (HPA), and 32 healthy controls ^13^ was re-analysed for blood L-phenylalanine and L-tyrosine levels. Measured patient intakes and metabolite values were also obtained from the published data. Methodological details, including sample preparation steps and instrument parameters, can be found in ^13^.

### Infant microbiome data

To simulate the impact of the microbiome on newborns with PKU, a subset of the ERP005989 dataset was used. This dataset was originally from a study by Bäckhed et al. ^36^, which contained extensive metagenomics data and metadata on the infant gut microbiome for healthy individuals. We selected a total of 48 female infants between 1 and 13 days of age, which had not received antibiotics. Samples were included regardless of delivery type (vaginal or caesarean) or feeding regimen (breastfed or formula-fed), ensuring a diverse, yet controlled representation of early infant microbiomes.

### Microbiome composition analysis

Alpha diversity of the 48 gut microbiome samples from healthy female infants was assessed using species richness and Pielou’s evenness, based on metagenomic data (**Table S4**). For each sample, both metrics were calculated to quantify the number of microbial species and the uniformity of their distribution. Summary statistics, including the mean, standard deviation, minimum, and maximum values, were reported (**Table S5**, 6). Microbial composition was analysed at the phylum and genus levels. For each taxon, relative abundances were calculated across all samples, and descriptive statistics were computed, including the number of samples in which each taxon was detected **(Tables S5–S6**). The stacked bar plot showing the phylum-level relative abundances across all samples was generated in MATLAB R2022b using a customised version of the bar function.

### Infant microbial community models

For the selected 48 infant microbiomes, we obtained the corresponding microbial community metabolic models, which have been published within the APOLLO resource ^33^ from https://dataverse.harvard.edu/dataverse/APOLLO. Briefly, these microbial community models have been assembled using the Microbiome Modelling Toolbox ^64^, which is part of the COBRA Toolbox v3 ^65^, using the strain information provided within the ERP005989 dataset and their relative abundance. Note that these microbial community models consist of individual microbial metabolic models, which have been combined into one microbial community model per sample while maintaining the integrity of each microbial model and allowing for metabolite exchange between the microbes. Hence, each microbial metabolic model contains its own biomass reaction that accounts for the biomass precursors required to create a new cell and an associated strain-specific biomass compound. Each metabolic reaction in a microbial metabolic model was coupled to its biomass reaction such that

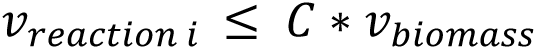

where C is a capacity constraint. For this study, we chose a capacity coefficient of 20,000 to identify maximal possible contributions of microbes to blood biomarker fluxes. The relative microbial abundance information in a given sample was used to set the stoichiometric coefficients of the respective strain in the microbial community biomass reaction, which sums all microbial biomass reactions. The microbial community biomass reaction was constrained to one, corresponding to one faecal emptying per day.

### Microbiome-associated infant-WBMs

Each of these microbial community models was put into the large intestinal lumen compartment of the female infant-WBM using the PSCM toolbox ^42^, which is part of the COBRA Toolbox v3 ^65^. By doing so, we enabled bidirectional exchange between the host model and the microbiome models. The resulting 48 microbiome-associated infant-WBMs only differed in their microbial composition.

### PAH-deficient infant-WBMs

To simulate PKU, we set the lower and upper bounds of the reactions associated with the PAH gene, i.e., PHETHPTOX2 and r0399, in all organs they appear to zero. This approach resulted in one germ-free PKU infant-WBM and 48 microbiome-associated PKU infant-WBMs, in addition to the aforementioned healthy counterparts.

### Prediction of PKU specific biomarkers using flux balance analysis

We predicted for all infant-WBMs the maximal possible flux value for 233 blood metabolites. The list of metabolites was selected based on being used in newborn screening programmes or reported to be biomarkers for IMDs in IEMBase ^34^. Subsequently, fluxes through these biomarkers in the blood compartment were computed using the function *performIEMAnalysis.m* from the COBRA Toolbox ^65^. Briefly, for the simulation of any IMD, the reactions *k* = (*k*_1_,…,*k_n_*)*^T^*associated with the defect gene in any organ were identified (i.e., PHETHPTOX2 and r0399). A dummy reaction *v_dummy_* was added to the WBM, summing the equal contribution of each reaction *k_i_*. The flux through *v_dummy_* was then maximised. The maximal possible value *z* for *v_dummy_* was used to set the lower bound on *v_dummy_* to 75% of this value. Afterward, an artificial demand reaction in the blood compartment *v_demand_* using the *addDemandReaction.m* function was added for each biomarker independently. This demand reaction was then used as the objective function for the flux maximisation in the linear programming (LP) problem. The corresponding LP problem (i.e., flux balance analysis) was:

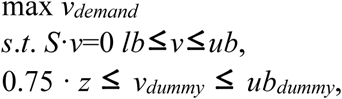

where *S* and *v* correspond to the stoichiometric matrix and flux vector, respectively, and reactions *v_dummy_* and *v_demand_* added to the original model. For the knock-out (i.e., PKU) infant-WBM, the lower and upper bound of *v_dummy_* was set to 0, and the maximisation through each biomarker metabolite demand reaction *v_demand_* was determined. Finally, the obtained flux values within the biomarker reaction of the wild-type (healthy) model *v_WT_* and the diseased model *v_D_* were compared. The relative flux increase was calculated as

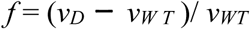

This approach allowed us to compare the predicted relative flux increase for the list of biomarkers. All units are given in mmol/infant/day if not stated differently.

### Shadow price analysis

To evaluate the impact of microbial activity on host metabolism under healthy and PKU conditions, we conducted a shadow price analysis. The shadow price is returned as dual to each primal linear programming problem solved, i.e., it is part of the flux balance solution returned by the COBRA Toolbox v3. The shadow price indicates by how much to objective value would change if the flux through the metabolite would increase or decrease by one unit:

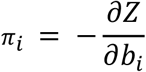

Where *π*_*i*_ represents the shadow price, Z is the objective value, and *b*_*i*_is the metabolite. As each microbial metabolic model in the microbial community model had its own strain-specific biomass compound, each microbe effectively corresponded to a “metabolite” and had an associated shadow price.

To identify the microbes that would most affect the maximal possible L-tyrosine blood flux value (or L-phenylalanine), we defined “strongly negative shadow prices” (SNSP). Therefore, we calculated Z-scores for microbe-associated shadow prices:

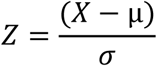

Where Z represents the Z-score of the data point, X is the shadow price value, μ is the mean of the dataset, and σ is the standard deviation of the dataset (Navidi, 2006). We then identified all those microbes that had a Z-score higher than 1 and classified them as SNSP.

### Flux taxon analysis

The relative abundances of microbes, stratified by sample sex, were correlated with tested biomarkers fluxes derived from host-microbiome model simulations. Statistically significant correlations, defined by p-values less than 0.05, are reported to highlight microbial taxa with a potential impact on L-tyrosine production.

## Code and data availability

All code is available at <u>COBRA Toolbox</u>. The toolbox installation procedure can be found at: https://opencobra.github.io/cobratoolbox/stable/installation.html. An MLX script reproducing the study step by step is available in the papers\2025_PKU folder.

All microbiome-associated infant-WBMs are available at: https://dataverse.harvard.edu/dataset.xhtml?persistentId=doi:10.7910/DVN/OUSN8Y Pre-run simulation results are available at: https://dataverse.harvard.edu/dataset.xhtml?persistentId=doi:10.7910/DVN/DW9UJE

All simulations were conducted using MATLAB 2020b (Mathworks, Inc.) as the programming environment, ILOG CPLEX (IBM, Inc.) as the LP/QP solver, and COBRA Toolbox v3.0 ^65^.

## Supporting information

Reproducibility Instructions

## Acknowledgements

We thank Jadzia Murphy for their help with metabolomic metadata and Bram Nap for their comments on the manuscript.

## Author contributions

IT and MM designed the study. JW performed taxonomic profiling of metagenomic data. MM performed simulations. LH and US contributed data. MM, TH, and IT analysed data. MM and IT drafted the manuscript. MM and FM developed the mlx code for reproducing the study results. All authors revised the manuscript. IT supervised the study. LH, US, and IT acquired funding.

## List of Supplementary Tables

- **Supplementary Table S1**: Flux simulation results for 233 biomarkers in germ-free whole-body models under healthy and PKU conditions.
- **Supplementary Table S2**: Detailed characteristics of the infant samples, including feeding regimen, mode of delivery, and sampling day.
- **Supplementary Table S3**: Relative abundance of bacterial phyla across female infant samples.
- **Supplementary Table S4**: Alpha diversity measures (species richness and Pielou’s evenness) for 48 gut microbiome samples.
- **Supplementary Table S5**: Mean, standard deviation, and distribution of bacterial phyla across infant gut microbiome samples.
- **Supplementary Table S6**: Mean, standard deviation, and distribution of bacterial genera across infant gut microbiome samples.
- **Supplementary Table S7**: Flux simulation results for 9 biomarkers in healthy and PKU microbiome samples.
- **Supplementary Table S8:** Relative reaction abundance for tyrosine production pathways in the 48 female infant metagenomic samples.

