## Supplementary material for "Elucidating a potential role of the infant gut microbiome on the bioavailability of L-tyrosine in phenylketonuria": Reproducibility Instructions

Steps to Reproduce the Study

1) Install COBRA Toolbox:
Follow the installation procedure outlined in the official documentation:
https://opencobra.github.io/cobratoolbox/stable/installation.html

2) Download the Infant Germ-Free and 48 Microbiome-Personalized WBMs:
Download the relevant models from the dataset available at:
Infant Models Used for PKU Analysis
https://dataverse.harvard.edu/dataset.xhtml?persistentId=doi:10.7910/DVN/OUSN8Y

3) Store the Models:
After downloading, place the infant germ-free models and the microbiome-personalized WBMs into the COBRA Toolbox remote directory:
papers\2025_PKU\HM_models\female\CC_20000.

4) Running the Full Simulation:
You can now test the study setup running the PKU_Biomarker_MicrobiomePipeline.mlx file.
Please note that executing the entire pipeline, particularly Step 3 of the .mlx file, may require several days of simulation time.

5) Testing Specific Steps:
Please be aware that Steps 4 and 5 require the full execution of the simulation of Step 3, to ensure proper functionality.
To save time, we have provided the pre-stored simulation results, which you can download and use as an alternative to running the full simulation.

5.1) Download the pre-stored results of the simulation from the following dataset:
Pre-Stored Results PKU Analysis
https://dataverse.harvard.edu/dataset.xhtml?persistentId=doi:10.7910/DVN/DW9UJE

5.2) After downloading, unzip the files and store the folders named according to the VMH IDs of metabolites in the directory:
papers\2025_PKU\solutionDir\Pre_stored_PKU.
